# Lethality of West Nile virus strains in a mouse model

**DOI:** 10.1101/2023.12.22.572994

**Authors:** A.E. Siniavin, I.V. Dolzhikova, A.A. Pochtovyi, D.A. Reshetnikov, V.A. Gushchin

**Author notes:** Corresponding author: Inna Dolzhikova.

## Abstract

West Nile fever (WNF) is a viral infection caused by West Nile virus (WNV), a flavivirus of the *Flaviviridae* family. Virus circulates between mosquitoes and wild birds, but can infect other species, including humans. The first cases of West Nile fever were reported in Africa in the 1930s. Currently, WNV has a wide geographic range, which includes countries in Europe, Asia, Africa, Australia, and North and South America, where it periodically causes WNF outbreaks. The disease in human occurs with the development of fever, and in some cases ending up severe neurological complications. Studies of the virus in animal models demonstrate that virulence varies depending on the host species, the genotype of the virus, and the presence of substitutions in key viral proteins, even within the same genotype. These studies highlight the need for comparative studies of different WNV strains to evaluate the impact of amino acid substitutions on WNV pathogenesis. Analysis of key mutations and substitutions will allow the development of a safe and effective vaccine for the prevention of WNF.

## Introduction

WNV occurs in people with the development of fever, which in some cases causes severe neurological complications (meningitis, encephalitis or acute flaccid paralysis). Fever develops in approximately 20-25% of patients, and the severity of the disease varies widely depending on the lineage of the virus. Damage to the nervous system can lead to the death of the patient; mortality rates during various outbreaks and epidemics range from 4 to 14%. The development of a neurological form of WNV can cause various lifelong complications in people who have suffered a severe form of the disease. There are currently no approved for use in human vaccines in the world to prevent West Nile fever. Vaccines are being developed in different countries. One of the key parameters assessed when developing vaccines is the ability to form protective immunity, which allows protection against infection. For this purpose, infectious animal models are used that are capable of simulating the infectious process and assessing the effectiveness of the developed tools of prevention and treatment. The most convenient to use are mouse models [1-2], but there are not many models for West Nile virus, especially for the relevant genetic variants. We conducted a study of the lethality in a mouse model of several strains of West Nile virus belonging to genotype 1, which is the most widespread throughout the world.

## Methods

### Cell and viruses

Vero E6 cells (ATCC #CCL-81) were grown in DMEM (Gibco, Carlsbad, CA) supplemented with 10% fetal bovine serum (FBS; Hyclone), 1× GlutaMAX, 1× Anti-Anti solution (all from Gibco, USA), and incubated at 37°C in 5% CO_2_. West Nile virus strains Eg101 (received in 1951), LEIV-1628Az (isolated in 1967), LEIV-1640Az (isolated in 1967) and Az5 (isolated in 1977) were prepared by inoculation a confluent monolayer of Vero E6 cells and a clarified harvest of the culture medium was collected after 5 days of incubation. Virus stocks were titrated by end-point dilution assay on Vero E6 cells. The virus titer was expressed as TCID_50_/ml.

### Sequencing

Viral RNA was fragmented, and the first cDNA chain was synthesized with random hexamer primers using the RevertAid First Strand cDNA Synthesis Kit (Thermo Fisher Scientific, Waltham, MA, USA), followed by the second DNA chain synthesis using NEBNext Ultra II Non-Directional RNA Second Strand Synthesis Module (New England Biolabs, Ipswich, MA, USA). The obtained cDNA was used to prepare a NEBNext DNA Library Prep Set for Ion Torrent (New England Biolabs, Ipswich, MA, USA) and barcoded with the use of Ion Code Barcode Adapters (Thermo Fisher Scientific, Waltham, MA, USA) according to the manufacturer’s instructions. DNA sequencing was performed using the Ion S5 XL System (Thermo Fisher Scientific, Waltham, MA, USA). The raw data was filtered by quality and length using vsearch v2.13.3 (DOI: 10.7717/peerj.2584). The trimmed reads were de novo assembled into contigs by the SPAdes v3.14.0 (DOI: 10.1089/cmb.2012.0021) with the —iontorrent key. The viruses were sequenced, and protein substitutions were analyzed relative to the reference genome strain Eg101 (GenBank: AF260968.1).

### Animal studies

All animal experiments were performed in strict accordance with the recommendations of the National Standard of the Russian Federation (GOST R 53434–2009; Principles of Good Laboratory Practice). Normal 8- to 10-week-old BALB/c mice were purchased from the nursery of laboratory animals Stolbovaya and kept in individually ventilated isolator cages (IsoCage N Bio-containment System, Tecniplast). Animals were intraperitoneally (IP) inoculated with each WNV strain (6 lg TCID_50_ per animal) and observed for morbidity and mortality for 16 days.

## Results

We used 4 strains of West Nile virus genotype 1. Analysis of amino acid substitutions revealed a significant number of substitutions in three strains LEIV-1628Az, LEIV-1640Az and Az5 in the NS4B protein in most cases with unknown effect on viral phenotype. It is important to note that in the strain Eg101, the P156S substitution of the glycoprotein E was detected, which can lead to an increase in the neurovirulence of the virus [3].

**Table 1.** List of amino acid substitutions in structural and nonstructural proteins of West Nile virus. Substitutions that lead to an increase in virulence are marked in red, and substitutions that lead to a decrease in virulence are marked in green.

| Protein | West Nile Virus Strain |  |  |  |
| --- | --- | --- | --- | --- |
|  | Eg101 | LEIV-1628Az | LEIV-1640Az | Az5 |
| prM |  |  |  | K9R |
| E | **P156S** | E273X<br>V488I | V488I | T76A<br>V488I |
| NS2B |  | V11A | V11A | V11A |
| NS3 |  | H87Y<br>I155V<br>**P249T** | H87Y<br>**P249T** | H87Y<br>I155V<br>**P249T** |
| NS4B |  | E22D<br>A23V<br>A175T<br>S201P<br>K215E<br>H230Y<br>L238P<br>H245X | E22D<br>A23V<br>A175T<br>S201P<br>K215E<br>H230Y<br>L238P | E22D<br>A23V<br>A175T<br>S201P<br>K215E<br>H230Y<br>L238P |
| NS5 |  | D813G | D813G | G525D<br>D813G |

An analysis of the survival rates after WNV challenge showed that the Eg101 virus strain, which contains the glycoprotein E P156S substitution, is lethal for animals: all infected animals showed signs of a neuroinvasive infection (paralysis), the animals died by 10 days after infection (Figure 1). At the same time, an analysis of survival after infection with strains 1628, 1640 and Az5 at same viral dose (6 lg TCID_50_) showed a mortality rate of 75-25%, and signs of neuroinvasive infection were detected only in animals that subsequently died.

**Figure 1.**
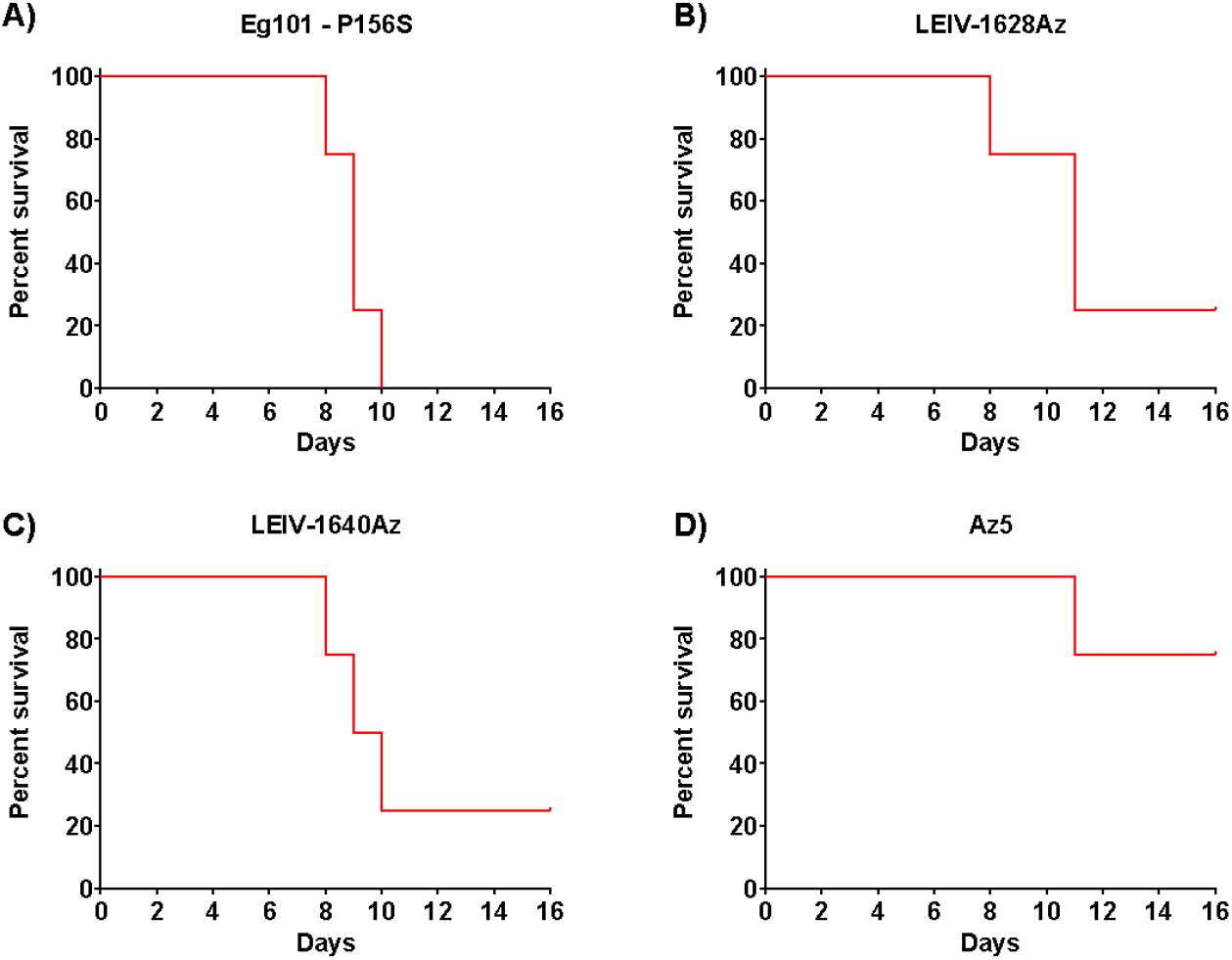
Survival rates of mice after infection with 6 lg TCID_50_ West Nile virus strains Eg101 (A), LEIV-1628Az (B), LEIV-1640Az (C), and Az5 (D) for 16 days.

## Discussion

WNV was first discovered in the West Nile province of Uganda in 1937 in a patient with a febrile illness [4–6]. To date, the virus has spread all continents except of Antarctica. It should be noted that the prevalence and severity of the disease varies greatly depending on the geographic area. For example, WNV is one of the leading causes of encephalitis in North America, and in Australia the disease in most cases occurs without neurological complications. Such differences may be caused by the evolutionary and geographically determined genetic diversity of the virus, based on which WNV was grouped into 9 genotypes [7-11]. The most studied and widespread are WNV genotypes 1 and 2, which are characterized by a high incidence of neurological diseases [12]. Genotype 1 covers countries in Africa, Asia, the Middle East, Europe, Australia and America. The wide variety of circulating West Nile virus strains results in varying degrees of infection severity. For example, studies have shown that the T249P substitution in the NS3 protein leads to increased virulence [13-16]. Another important factor determining pathogenicity is the glycosylation of the envelope protein E [16-18]. In our study, we found that the West Nile virus strain Eg101, for which no lethal animal model has been described in the literature [19], contains the P156S substitution in glycoprotein E and leads to the death of 100% of animals after infection.

Although some cases of WNV infection may be asymptomatic in humans, ∼20% of those infected develop a febrile influenza-like illness with symptoms similar to those of dengue fever; ∼1 in 150 infected people are diagnosed with life-threatening neuroinvasive forms of the disease with the development of meningitis, encephalitis and acute flaccid paralysis. Mortality rates for CNS lesions can reach 14% [20–22]. To date, there is no specific treatment for West Nile fever. There are currently no registered vaccines to prevent West Nile fever in humans. Active research is being conducted all over the world: in Europe, the USA, Russia and other countries. Several vaccine candidates based on different technology platforms (inactivated, chimeric, subunit, DNA, and adenoviral vector vaccines) are currently in various stages of preclinical and clinical studies [23– 26]. In vaccine development, a key parameter assessed in preclinical studies is the ability of the vaccine to generate protective immunity that protects against West Nile virus infection. Several animal models have now been described, with the mouse model being the most common. Studies of the virus in animal models demonstrate that virulence varies depending on the host species, the genotype of the virus, and the presence of substitutions in key viral proteins, even within the same genotype. These studies highlight the need for comparative studies of different WNV strains to evaluate the impact of mutations on WNV pathogenesis. Analysis of key mutations and substitutions will allow the development of a safe and effective vaccine for the prevention of WNF.

